# Morphological Plasticity and Reproductive Strategies of *Kalanchoe* Species in Invasive Spread

**DOI:** 10.1101/2024.12.25.630291

**Authors:** Zhe Zhang, Daisuke Sugiura, Wataru Yamori, Yanhong Tang

## Abstract

*Kalanchoe* species, originally introduced worldwide as ornamentals, are now reported to be spreading in many regions, including China. It is hypothesized that morphological plasticity and the production of asexual plantlets contribute to their successful invasion. To address the hypothesis, four species - *Kalanchoe delagoensis* Eckl. & Zeyh., *Kalanchoe × houghtonii* D. B. Ward, *Kalanchoe laetivirens* Desc. and *Kalanchoe daigremontiana* Raym. - Hamet & H. Perrier - were cultivated under contrasting light and water conditions, and their reproductive and vegetative traits were examined. The plants subjected to high light tended to prioritize plantlet production, accompanied by a reduction in vegetative growth. Two distinct reproductive strategies were observed: *K. delagoensis* and *K. × houghtonii* significantly increased plantlet production under high light conditions, whereas *K. daigremontiana* and *K. laetivirens* enhanced the fresh weight of individual plantlets without altering the total number produced. These results highlight the high plasticity of vegetative and reproductive growth in response to light and water availability. The increased production of plantlets may contribute to the widespread occurrence of *Kalanchoe* species in open fields.

## 1. Introduction

The term ‘pseudovivipary’ refers to the production of apomictic or asexual propagules, such as bulbils or plantlets, enabling plants to reproduce asexually. The phenomenon is most commonly observed in plants that inhabit arctic, alpine or arid environments, where they are subjected to prolonged periods of stress (Elmqvist and Cox 1996). In such habitats, the probability of offsprings dispersing to more favorable locations in space or time is often very low, rendering the production of dormant seeds less advantageous. Pseudovivipary offers specific benefits in these conditions, as the plantlets continue to grow and derive nutrition from the maternal parent until detachment. This allows the detached young plantlets to develop enhanced survival abilities in harsh environments.

Some species in the genus *Kalanchoe* (Crassulaceae) are capable of spontaneously developing plantlets along the notches of their leaf margins in addition to reproducing sexually (Howe 1931). Two types of pseudovivipary exists in *Kalanchoe* genus; some species form plantlets only under stress (induced plantlet-forming species), while others produce plantlets spontaneously (constitutive plantlet-forming species). The latter phenomenon has been attributed to the genetic differences in regulation of embryogenesis (Garcês et al. 2007, Jacome-Blasquez and Kim 2023). Additionally, they all exhibit the characteristics of succulent CAM (Crassulacean acid metabolism) plants. The leaves of many *Kalanchoe* species are capable of storing water in mesophyll cells, thereby enabling the plant to become temporarily independent from external water supply and to retain physiological activity against short-term drought (Eggli and Nyffeler 2009, Perez-Lopez et al. 2023). Combined with their high water-use efficiency of CAM photosynthesis, these traits provide *Kalanchoe* species with strong competitive and colonizing abilities in tropical and subtropical arid environments.

*Kalanchoe* species are native to Madagascar and tropical Africa, inhabit predominantly in the arid to mesic savanna areas with summer-rainfall (Smith et al. 2019). Introduced worldwide as ornamental plants, the invasion of *Kalanchoe* species is now reported in many regions, threatening local biodiversity. Research in Venezuela showed that the native seedling recruitment is negatively affected by the invasion of *K. daigremontiana*, which significantly lowered the density of native seedlings and species richness of native vegetation (Herrera et al. 2016). The suppression of native plant growth may be attributed to the presence of allelochemicals, as evidenced by the inhibition of seed germination and seedling development observed in plants grown with *K. daigremontiana* and its extracts (Groner 1975).

The primary trait contributing to the invasive success of *Kalanchoe* species is likely the rapid growth and dispersal of asexual propagules. Population modeling has revealed that the establishment of *K. daigremontiana* depends exclusively on plantlet recruitment (Herrera et al. 2012). Consequently, eradication through plant removal can only be achieved if initiated shortly after introduction, as pseudovivipary enables rapid population growth during the initial phases of invasion. Asexual plantlets showed high survival rate (75-100%) in the field, and seeds are also present in the seed bank by autogamous despite low viability and germination rates (Herrera and Nassar 2009). This led to the potential for invasion of Neotropical arid zones mainly via asexual reproduction and dispersal. The estimation of genetic diversity of *K. delagoensis, K. daigremontiana* and their artificial hybrid *K. × houghtonii* (Gideon 2020) in Mexico showed low levels of genotypic diversity, indicating that the invasion of these species occurred from the introduction of a single genotype and has expanded primarily through clonal growth and human-mediated dispersal. Furthermore, the occurrence of natural hybridization is unlikely in the wild (Guerra-García et al. 2015).

The issue of *Kalanchoe* invasion is becoming increasingly prevalent worldwide in recent years. Although only *Kalanchoe pinnata* (Lam.) Pers., which exhibits plantlet formation in response to stress, is included in the list of invasive alien species (Xu et al. 2012), other constitutive plantlet-forming species have been identified and are spreading rapidly in the north of China as well, such as *K. delagoensis, K. daigremontiana, K. × houghtonii* (Wang et al. 2016), and *K. laetivirens*, which has received comparatively less attention (Fig. S1). Initially introduced as ornamental plants, *Kalanchoe* species are primarily distributed in urban and suburban areas. Their spread occurs mainly through clonal propagation, which limits long-distance dispersal. However, the expansion of these species between cities is largely attributed to human activities, including garbage disposal, online shopping, mailing, and other similar practices.

The global distribution of these species, including their occurrence in China, exhibits a similar pattern. All four species have been observed to acclimate well to urban ecosystems and reproduce via plantlets regardless of light and water conditions in tropical and subtropical areas. However, their distribution appears to be constrained primarily by the temperature of the coldest month. Among these species, *K. delagoensis* and *K. × houghtonii* have been observed with much greater frequency and higher abundance compared to *K. daigremontiana* and *K. laetivirens*. Although the reliability of occurrence data may be influenced by the time of introduction and the level of awareness among local citizens, *K. delagoensis* and *K. × houghtonii* seem to possess a stronger ability to escape cultivation, acclimate broadly, and spread extensively in urban environments.

There is currently a lack of comprehensive knowledge regarding the ecophysiology of *Kalanchoe* and the scientific evaluation of its invasiveness. It is therefore important to gain a deeper understanding of how *Kalanchoe* pseudoviviparous plants acclimate to the urban ecosystems worldwide, how they gain competitive advantages through asexual reproduction in the field and what characteristics lead to the difference in abundance. The aim of this study is to elucidate the interactive effect of light intensity and water availability on the vegetative growth and asexual reproduction of *Kalanchoe* plants and to investigate how they balance the growth and reproduction individually. Four constitutive plantlet-forming species that are commonly found wild in China were selected and cultivated in contrasting light and water conditions. Their plasticity and acclimation strategies were analyzed by comparing morphological, physiological and reproductive traits.

## 2. Materials and Methods

### 2.1 Plant materials and growth condition

The plants of *K. delagoensis, K*.*× houghtonii, K. daigremontiana* and *K. laetivirens* were obtained from a roof garden in Jinan city, Shandong Province, China. The plants were observed to be between 10 and 15 cm in height. The light conditions were those of natural sunlight, with slight shading. The midday mean photosynthetic photon flux density (PPFD) on a sunny summer day was 673 μmol m^-2^ s^-1^. In addition to natural precipitation, the plants were watered every 3 days throughout the summer period. Subsequently, the plants were randomly assigned to four treatment groups and cultivated in a laboratory environment with a temperature of 25 °C and a 12-hour light interval for a period of 6 months. The plants in the high light and dry (HD) group were watered 100 ml once a week and grown under 800 μmol m^-2^ s^-1^ PPFD. The plants in the high light and wet (HW) group were watered 100 ml twice a week and grown under a PPFD of 800 μmol m^-2^ s^-1^. The plants in the low light and dry (LD) group were watered 100 ml once in two weeks and were grown under a PPFD of 60 μmol m^-2^ s^-1^. The plants under the low light and wet (LW) group were watered 100 ml once a week and cultivated under 60 μmol m^-2^ s^-1^ PPFD. The minimum soil water content for the dry groups was 5 %, while the wet groups had a minimum of 30 %. To avoid the impact of LED light heating on air temperature from affecting plant growth, fans were installed to enhance air circulation.

### 2.2 Sampling and measurements

For each species, three individuals were randomly selected and assigned to each treatment group for measurement. New leaves and old leaves were collected from each individual to analyze the effect of leaf developmental stage on acclimation strategy and the plasticity of leaf traits. In the case of *K. delagoensis* and *K*.*× houghtonii*, the leaves that germinated and matured in the treatment environment were considered to be the new leaves. The 5^th^ pair of leaves were sampled. The leaves that matured in the primary environment but did not senesce in the treatment environment were considered as the old leaves. The 15^th^ pair of *K. delagoensis* leaves and the 10^th^ pair of *K. × houghtonii* leaves were sampled. In the case of *K. daigremontiana* and *K. laetivirens*, as the old leaves had already reached a state of senescence before the new leaves had matured in the treatment environment, the leaves that germinated in the treatment environment but had not reached full maturity were considered as the new leaves, with the 3^rd^ pair of leaves sampled. The leaves that germinated in the primal environment and had matured in the treatment environment were considered to be the old leaves, with the 5^th^ pair of leaves sampled.

#### Individual morphology

The number of leaves was counted directly. The fresh weight of the shoot and root were measured separately after cutting stem from the soil surface, and the root/shoot ratio was calculated by dividing the root fresh weight by the shoot fresh weight. The stem length was measured before and after treatment, and their difference value was determined as stem elongation. The stem diameter was measured with ImageJ by measuring the diameter of stem cross section under a microscope.

#### Asexual reproduction

During the cultivation, some plantlets fell off spontaneously, thereby precluding the possibility of measuring the quantity and quality of plantlets directly at one time. At the end of treatment, the remaining plantlets were collected, counted and the total mass was measured. The fresh weight of each plantlet was calculated by dividing the total mass measured by the number of plantlets included in the measurement. The number of plantlets was obtained by counting the abscission scar of plantlet detachment on the leaf margin. The total mass of the plantlets was calculated by multiplying the fresh weight of each plantlet by the quantity of plantlets. The reproduction ratio was calculated using the following equation below to evaluate the proportion of biomass investment allocated to vegetative growth and asexual reproduction.

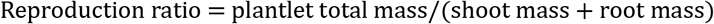

#### Leaf morphology

The leaves were scanned and the dimensions of length, width and area were then determined using the image analysis software ImageJ. Leaf thickness was measured with the ImageJ using the photos of fresh transection of midrib captured under microscope. Specific leaf area (SLA) was calculated by dividing leaf area with leaf dry weight. The upper and lower epidermis were torn with forceps directly, and the length of stomata was measured using images of stomata under a microscope with ImageJ. Stomatal density was calculated using epidermal impressions made with clear nail polish.

#### Leaf physiology

Leaf fresh weight was measured immediately after detachment. The saturated fresh weight was measured after the leaves were soaked completely in distilled water for 24 hours. The leaf dry weight was subsequently obtained after the leaves had been dried in an oven at 65 °C to constant weight. The leaf saturated water content (SWC) and relative water content (RWC) were calculated with following equations (Ogburn and Edwards 2012).

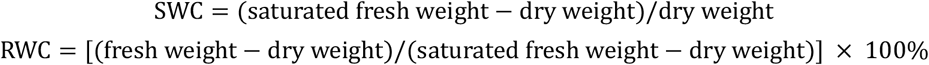

The content of photosynthetic pigments was quantified using a spectrophotometric method. 0.4 g of each fresh leaf without midrib was cut into 2 mm wide short pieces and soaked in 10 ml absolute ethyl alcohol within centrifuge tube, then extracted at 4 °C for 48 hours in the dark until the tissues turned white. The absorbance at wavelengths of 665 nm (A_665_), 649 nm (A_649_) and 470 nm (A_470_) were determined using an ultraviolet spectrophotometer, and the contents of photosynthetic pigments were calculated with following equations (Lichtenthaler 1987).

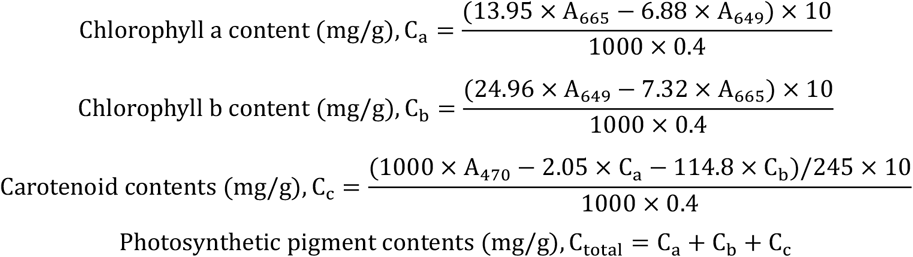

### 2.3 Statistical analysis

A one-way analysis of variance (ANOVA) and Duncan’s multiple comparison test were performed to determine the existence of differences and the significance of these differences in trait means among treatments with a 95% confidence interval. A two-way ANOVA was conducted to examine the effects of light, water and their interactions on plant traits. Pearson correlation analysis was conducted to reveal the mutual relationship between traits at the individual and leaf levels. Principal component analysis was conducted with the Origin 2022 app Principal Component Analysis v1.50. Linear and non-linear fitting were conducted with Origin 2022. To compare the plasticity among traits and among species, the plasticity index (PI) for each trait was calculated using Valladares’s equation (Valladares et al. 2000).

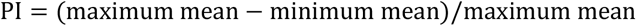

## 3. Results

### 3.1 Vegetative growth of pseudoviviparous *Kalanchoe* species

Under the conditions of low light intensity, a significant elongation of the stems was observed, and a notable difference existed between the water conditions as well, with the plants grown in LW exhibited the greater stem elongation (Fig. 1b, c). In species with bigger leaves (i.e. *K. daigremontiana* and *K. laetivirens*), greater leaf longevity and leaf number were observed under the low light environments, whereas leaves exhibited earlier senescence and detachment under the high light conditions. (Fig. 1a, b, Fig. S2). However, no such difference was observed in the case of *K. delagoensis* and *K*.*× houghtonii*. Additionally, there was little variation in the stem diameter across treatments (Fig. S2). Shoot fresh weight was higher in low light environments, while root fresh weight exhibited the opposite trend (Fig. S2). Root/shoot ratio was approximately twice as high in low light conditions under high light, and differences under the water conditions were observed for *K. delagoensis* and *K*.*× houghtonii*, with a higher root/shoot ratio in the LD treatment (Fig. 1d).

**Fig. 1.**
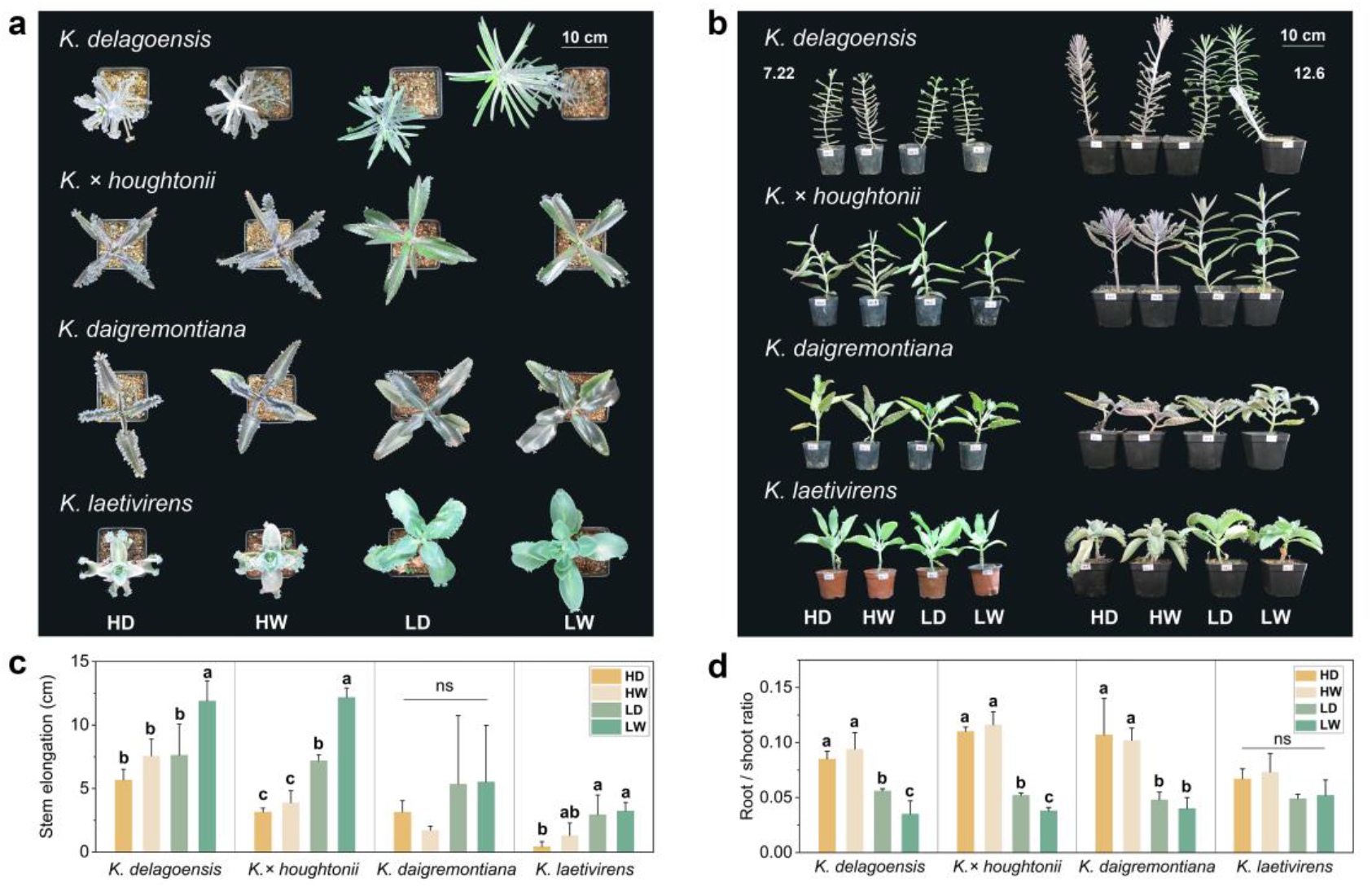
Individual morphology and biomass allocation of four *Kalanchoe* species under various environmental conditions. (a) The top view of plants grown in various environmental conditions. The plants grown in low light tend to have horizontal flat leaf and increased projection area. (b) The front view of plants from primal environment (left, July 22th) and grown in treatment environments (right, Dec. 6th). (c) Stem elongation of each species under each treatment. (d) Root/shoot ratio of each species under each treatment. Difference of alphabet above the error bar shows significance, while *ns* means no significant difference between the treatments

Leaf morphology varies among species (Fig. 2a). The leaves of *K. delagoensis* are narrowly subcylindrical in shape, displaying reddish-brown spots and grooves on the adaxial surface. Additionally, three to nine conical teeth are present at the apex, where plantlets emerge between the teeth. In low-light environments, leaves exhibited lengthening and thinning, with minimal change in width (Fig. 2b, Fig S3). New leaves and old leaves both showed morphological plasticity to adjust the leaf shape to acclimate to the environment. The leaves of *K*.*× houghtonii* are boat-shaped with serrated margins and purple splotches beneath. New leaves became longer and wider with lower thickness under low light, while old leaves showed almost no morphological change among treatments (Fig. 2b, Fig S3). The leaves of *K. daigremontiana* are triangular to lanceolate in shape, with a purple-blotched abaxial surface, serrate margins and an acute apex. There was a tendency for leaf length and width to increase in low light environment, which resulted in a change in leaf area (Fig. 2b, Fig S3). The new leaves grown in high light were thicker, but the old leaves showed no change in thickness. *K. laetivirens* has oblong to elliptic large bright green leaves. The leaf area of *K. laetivirens* showed no significant difference among treatments, indicating low morphological plasticity. However, leaves under high light curled and folded to reduce the area that exposed directly to light (Fig. 2ab, Fig S3). SLA showed a significant increase under low light conditions for all species (Fig. 2b).

**Fig. 2.**
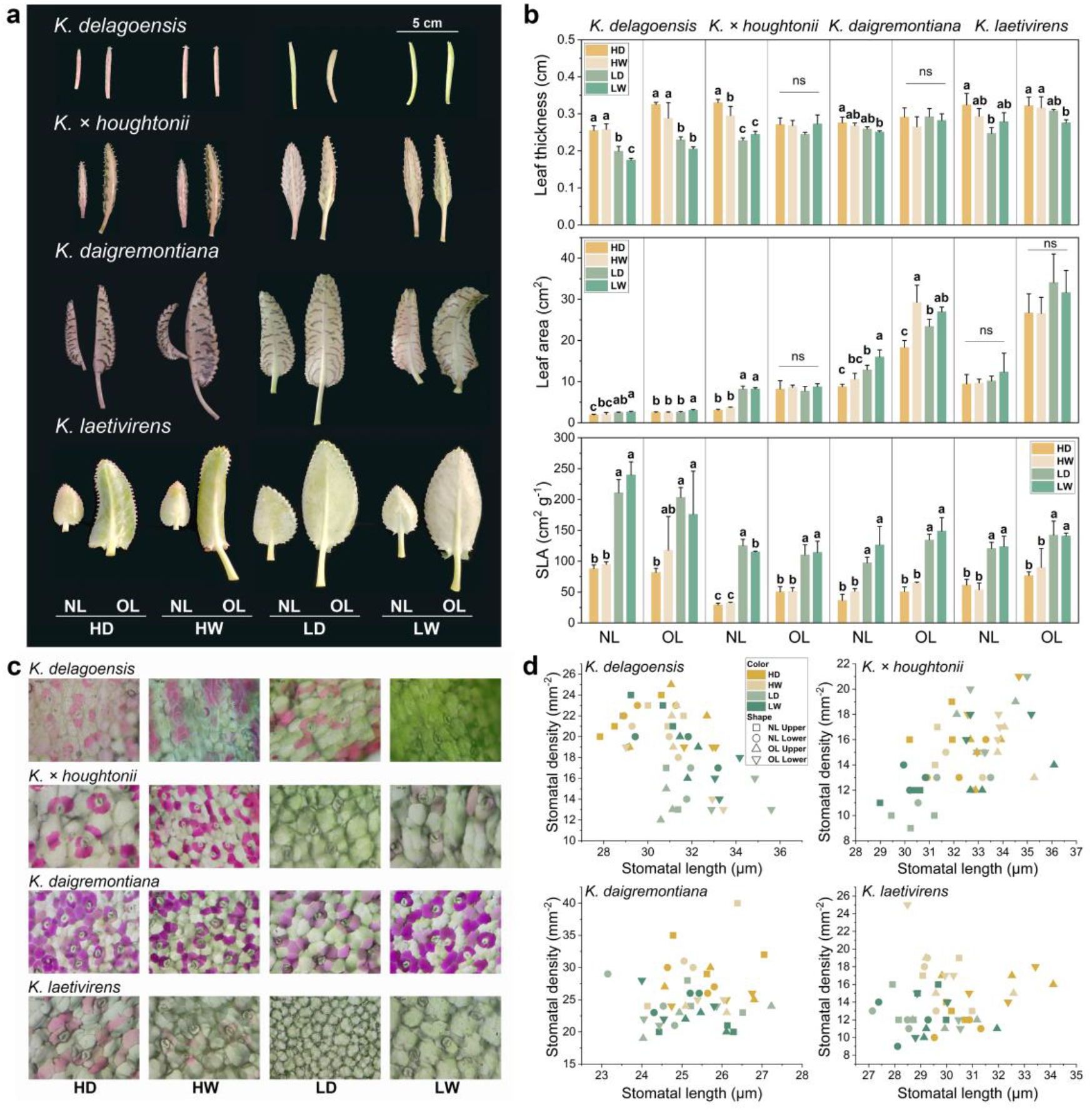
Leaf morphological traits of four *Kalanchoe* species under various environmental conditions. NL represents new leaves and OL represents old leaves. (a) Morphology of leaves under treatment environments. (b) Bar graphs of leaf morphological traits. Difference of alphabet above the error bar shows significance, while ns means no significant difference between the treatments. (c) Microscopic images of upper epidermis of leaf grown in various environmental conditions. Except for *K. daigremontiana* who shows no significant difference, the plants from high light or dry environments have more idioblast in leaf upper epidermis. (d) Scatter graphs of leaf stomatal length and density

The stomatal length of the four *Kalanchoe* species were highly conserved among treatments, with *K. daigremontiana* being around 23-28 μm and the other species about 27-37 μm (Fig. 2d, Fig S3). Leaves of *K. laetivirens* tended to have larger stomata under high light. Stomatal density exhibited variation, but no general tendency was observed. Most leaves tended to have more stomata under high light. *K. daigremontiana* had the highest stomatal density among the four species. An increase in the number of purple idioblasts in the upper epidermis under high light or dry environments was observed in *K. delagoensis, K*.*× houghtonii* and *K. laetivirens*, while upper epidermis of *K. daigremontiana* contains numerous idioblasts regardless of water and light conditions (Fig. 2c). The purple or pink substance in the idioblasts may be anthocyanin, which contributes to the stress resistance such as photoprotection (Steyn et al. 2002, Li and Ahammed 2023).

Leaves grown under high light accumulated significantly more dry matter than those grown in low light (Fig. 3a). However, leaves grown under low light had higher SWC, which was used to quantify succulence in succulent leaves (Ogburn and Edwards 2012), indicating the ability to store more water for the same amount of dry matter (Fig. 3b). The contents of chlorophyll a, chlorophyll b and carotenoids were significantly higher in leaves grown in low light, which may compensate for the loss of photosynthesis caused by the lack of light (Fig. 3c).

**Fig. 3.**
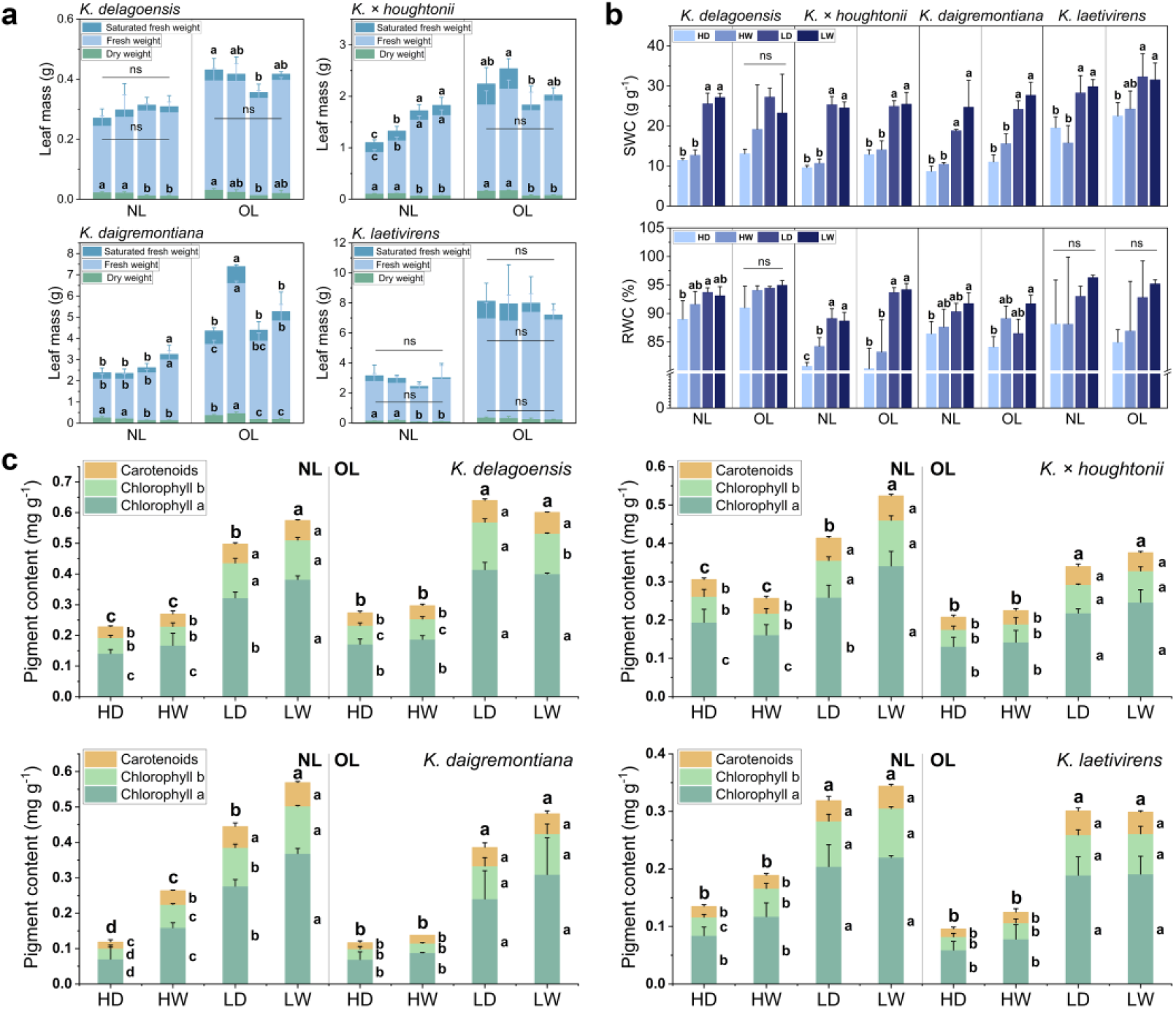
Leaf physiological characteristics of four *Kalanchoe* species under different environmental conditions. NL represents new leaves and OL represents old leaves. (a) Bar graphs of leaf dry weight, fresh weight and saturated fresh weight overlapping. (b) Bar graphs of leaf SWC and RWC. (c) Stacked bar graphs of leaf photosynthetic pigment contents. The alphabets at the top show the significance of total photosynthetic pigment contents, while the alphabets on the side of the bars show the significance of the individual pigment contents. The difference of the alphabet above the error bar shows the significance, while *ns* means no significant difference between the treatments

### 3.2 Asexual reproduction under varying light and water environments

As a possible main contributor of invasion, plantlet characteristics are of great importance for the survival and expansion of *Kalanchoe* populations. For *K. delagoensis* and *K*.*× houghtonii*, the number of plantlets was significantly higher in high light condition (Fig. 4d, Table S1). Fresh weight of individual plantlet was higher under low light for *K. delagoensis* especially in dry environment, while *K*.*× houghtonii* showed no such tendency. Both species exhibited no difference in plantlet total mass among treatments (Fig. 4e). The highest reproduction ratio was observed for *K. delagoensis* in LD and *K*.*× houghtonii* in HW. For *K. daigremontiana* and *K. laetivirens*, the number of plantlets produced showed no difference among treatments, but the fresh weight of individual plantlets exhibited significant increase under high light which also increased plantlet total mass, especially in *K. laetivirens* (Fig. 4d, e, Table S1). The reproduction ratio was highly increased in both species in high light condition. Plantlets germinated in low light and wet environments tended to have larger leaves, which is consistent with the morphological change of the maternal parent and gives them the ability to better acclimate to the same environment (Fig. 4a, b).

**Fig. 4.**
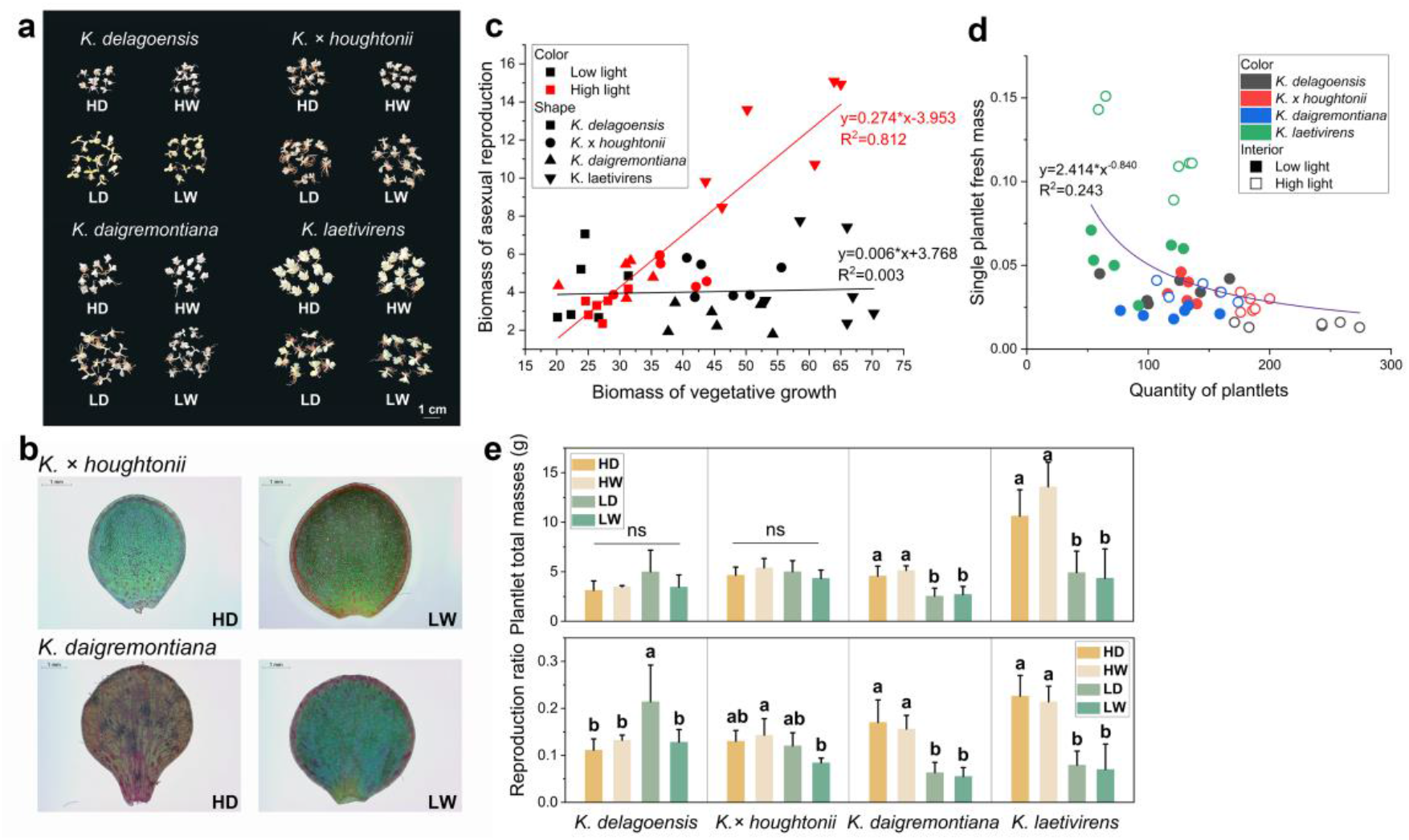
Asexual reproduction of four *Kalanchoe* species under different environmental conditions. (a) Morphology of plantlets produced under different environmental conditions. Ten plantlets of each species were showed for each treatment. (b) Single leaf of plantlet under HD and LW conditions. The plantlets under LW environment tend to have larger leaves. (c) Scatter graphs of vegetative growth biomass and asexual reproduction biomass under different light conditions (fresh). (d) Scatter graphs of fresh weight of individual plantlets and number of plantlets for each individual under different light conditions. (e) Bar graphs of asexual reproduction traits. Difference of alphabet above the error bar shows significance, while *ns* means no significant difference between the treatments

Four species showed little difference in plantlet number and fresh weight under low light (Fig. 4d). However, two types of strategy were observed under high light, as to produce more plantlets or to produce plantlets containing more nutrients (Fig. 4e). *K*.*× houghtonii* and *K. daigremontiana* were more conserved for reproductive traits among treatments, while *K. delagoensis* and *K. laetivirens* showed a strong response of reproductive traits for light conditions in two different directions (Fig. 4d).

### 3.3 Plasticity of four *Kalanchoe* species to light and water conditions

For each trait, the mean value of four species was calculated as the PI to investigate the acclimation patterns of pseudoviviparous *Kalanchoe* plants to light and water conditions. Stem elongation showed the highest plasticity among all traits that exceeds 0.7, followed by chlorophyll content (Fig. 5a). The length of stomata of the upper and lower epidermis, for which the PI was less than 0.06, were the most conserved traits among the traits tested in this research. Divided into five categories, leaf physiological traits exhibited high plasticity towards the change of light and water environments, especially for photosynthetic pigment content and SWC. Reproduction ratio had the highest plasticity in asexual reproduction traits and was similar to the root/shoot ratio, indicating that plants can flexibly adjust the allocation of resources on root growth, shoot growth and asexual reproduction to balance the growth and acclimate to the environments. The plasticity of plantlet mass was found to be higher than that of plantlet number. Leaf morphological traits showed low plasticity except for SLA. Among the highly conserved stomatal traits, the stomatal density of the upper epidermis (0.31) was observed to display a relatively high degree of plasticity.

**Fig. 5.**
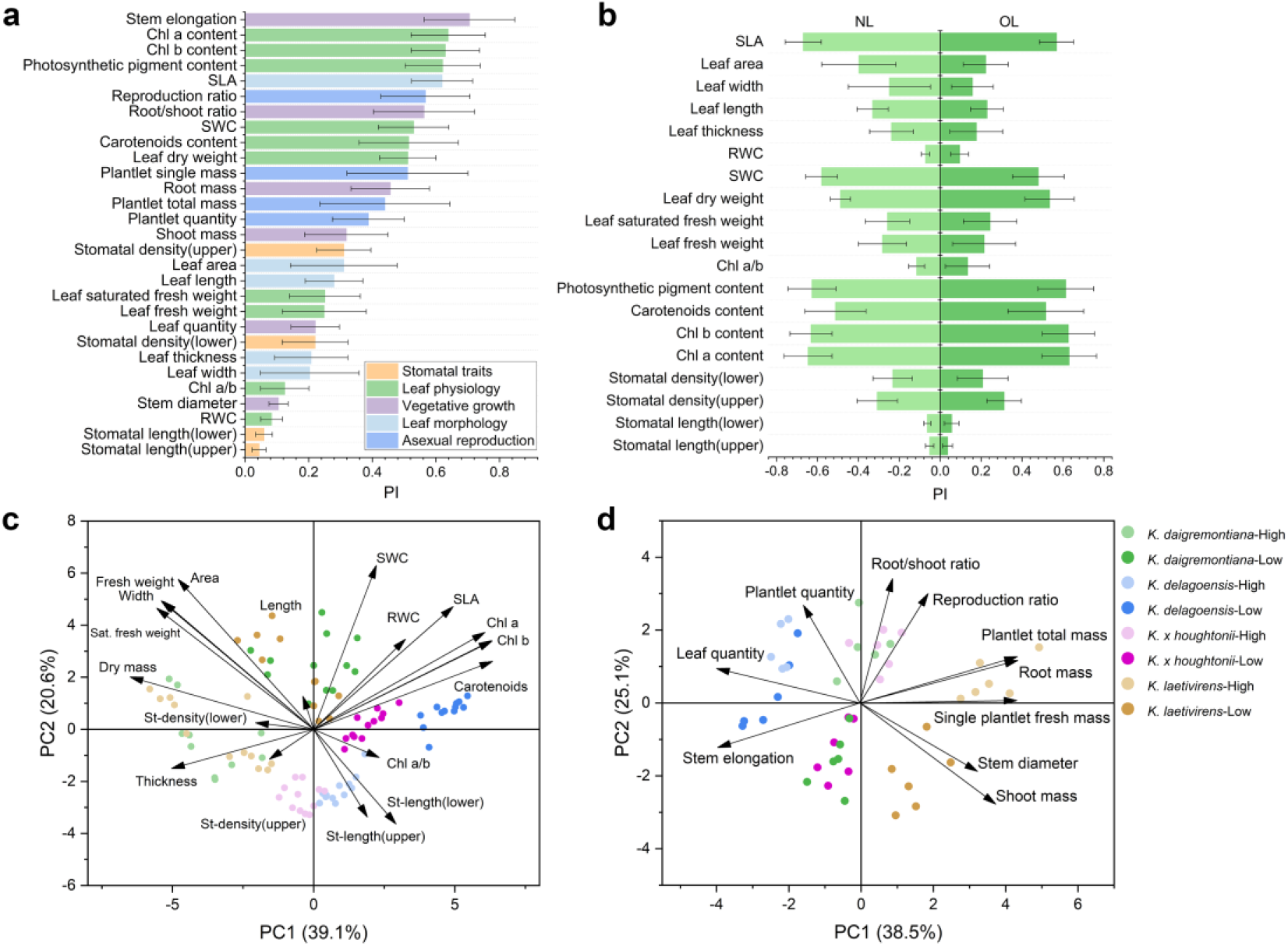
Plasticity and correlation of traits. Plasticity of (a) all traits and (b) new leaves (NL) and old leaves (OL) in comparison. PCA biplot of (c) leaf traits and (d) individual traits, showing different species and light intensity

For most leaf traits, new leaves showed greater plasticity than old leaves, especially for morphological traits, as old leaves had already reached a specific developmental stage or had even matured in both *K. delagoensis* and *K*.*× houghtonii* (Fig. 5b). However, despite this, old leaves showed some capacity to alter their shape in response to environment changes. Leaf physiological traits showed similar plasticity for new leaves and old leaves, suggesting that all leaves have strong capacity to physically adjust the metabolism to acclimate to their environment.

### 3.4 Correlation between traits

At the leaf level, the first two principal components collectively explained 59.7% of the observed variation (Fig. 5c). PC1 was found to be strongly correlated with chlorophyll a, b content, carotenoids content and dry mass, which may be indicative of photosynthetic capacity. Leaves that developed under low light increased the contents of photosynthetic pigments, while leaves that developed in high light showed an increase in the assimilation and accumulation of dry matter. PC2 was strongly correlated with fresh weight, saturated water content, and leaf area, which may reflect the water-storage capacity. Leaves that developed in low light conditions showed an enhanced capacity to store water. *K. daigremontiana* and *K. laetivirens* showed a higher similarity with regard to leaf traits.

At the individual level, the first two principal components collectively explained 63.6% of the observed variation (Fig. 5d). As a hybrid of *K. delagoensis* and *K. daigremontiana*, individual of *K. × houghtonii* appeared to show a higher degree of similarity to *K. daigremontiana*. The stem elongation, leaf number and fresh weight of individual plantlet mainly reflect the strategic difference among species, with *K. delagoensis* and *K. laetivirens* being the most distinct. The light intensity had a significant impact on the root/shoot ratio and reproduction ratio, suggesting that biomass allocation is highly flexible in response to changes in light environments. *K. delagoensis* was the only species that showed the strongest acclimation to water availability under low light conditions.

A strong negative correlation was observed between stem elongation and plantlet total mass, suggesting a potential trade-off between vegetative growth and asexual reproduction in response to changes of light environments (Fig. 5d, Fig. S4).

## 4. Discussion

### Acclimation of vegetative growth of *Kalanchoe* pseudoviviparous plants

The morphological and physiological changes observed in vegetative growth in response to light intensity were significant, with notable effects on stem elongation (Fig. 1c), root/shoot ratio (Fig. 1d), leaf shape (Fig. 2a, b) and photosynthetic pigment contents (Fig. 3c). Plants are capable of acclimating to low light environment through both shade-tolerance and shade-avoidance performance (Franklin 2008). The thinner leaves (Fig. 2b) and elevated chlorophyll contents (Fig. 3c) under low light condition observed in our study are likely to favor photosynthetic carbon gain, while the elongation of stems (Fig. 1c) is an acclimation trait to enable the plant to obtain more light resource and to overtop competing vegetations. Under the low light conditions, the stem of *K. delagoensis* exhibited a tendency to bend and crawl along the ground after reaching a certain height. This behavior may facilitate light capture in natural environments, potentially in conjunction with the phototropism (Liscum et al. 2014). It is pertinent to consider the vegetation of Madagascar, where these species originated. They evolved in an environment devoid of dense canopies and generally characterized by abundant sunlight and minimal dense canopy coverage. In these habitats, shading primarily arises from individuals of the same or similar species, which can reach heights of up to one meter and grow in close proximity (Eggli 2003, Rojas-Sandoval and Acevedo-Rodríguez 2022b, a). Under the high light conditions, plant size was reduced by the development of smaller leaves or changes in their angles to avoid capturing excessive light and overheating (Fig. 1a, 2a). The reddish coloration of the leaves is probably due to the anthocyanin accumulation in idioblasts beneath the epidermis (Fig. S3). This is supported by the evidence indicating a correlation between elevated anthocyanin concentrations and reduced photoinhibition (Li et al. 2015).

In *Kalanchoe* species with the capacity to form leaf plantlets, the leaves are not only responsible for carbon assimilation, but also for the storage of succulent water and asexual reproduction. The resource allocation among different organs is a critical factor in determining an organism’s life strategy (Weiner 2004). The optimal allocation theory posits that plants should allocate resources in a manner that increases their uptake of the resource that is most limiting to growth (Bloom et al. 1985). In high light environments, where water acquisition is often the primary objective, plants allocate more resources to the roots, resulting in an increased root/shoot ratio (Fig. 1d). Conversely, in low light environments, especially in wet environment with sufficient water, light becomes a limiting factor. This prompts the allocation of more resources to the aboveground organs to maximize light capture, leading to a reduction in root/shoot ratio (Fig. 1d).

### Morphological and structural plasticity in CAM plants

The importance of phenotypic plasticity in changing climate is widely reviewed in recent years as climate change has been shown to alter the availability of resources and the conditions that are crucial to plant performance (Matesanz et al. 2010, Nicotra et al. 2010, Stotz et al. 2021). CAM photosynthesis is highly plastic, intimately linked with the environmental factors and susceptible to changes in temperature, light intensity and water availability (Dodd et al. 2002). The metabolic basis and ecological significance of photosynthetic plasticity were studied intensively (Borland et al. 2011).

The morphological and structural plasticity of CAM plants is generally regarded as more constrained compared to the high plasticity of CAM photosynthesis, as physiological and biochemical plasticity appears to be more responsive and energy efficient (Luttge 2010). Nevertheless, this study revealed significant morphological plasticity in all the four species, ranging from individual leaves to entire plants (Fig. 5). This suggests that long-term or significant alternations in light and water environments can induce acclimation at both the physiological and morphological levels. Furthermore, succulents that originated from low heterogeneity environments, which are often perceived as less efficient in morphological changes than other plants (Alpert and Simms 2002), can actively modify their structure to acclimate to their environment. A concurrent assessment of physiological and morphological plasticity over time will facilitate a more comprehensive understanding of the remarkable acclimation strategy of *Kalanchoe* species to changing environments. This will include the measurement of physiological plasticity, such as photosynthetic characteristics and water status, as well as morphological plasticity including leaf shape, biomass allocation, and stem elongation.

The morphological and structural changes observed in this study were predominantly affected by light intensity rather than by water availability or their interaction (Table S2, S3). The relative stability of water status in succulent leaves may contribute to this effect. These leaves utilize stored water to prevent the development of low water potentials in their photosynthetic tissues, allowing them to remain independent of external water supply for extended periods (Ogburn and Edwards 2010). However, it should be noted that only two levels of water availability were tested in this study, which may have masked the full extent of these species’ acclimation capacity towards water limitations. To gain a more comprehensive understanding of this phenomenon, future studies should incorporate a broader range of more extreme water treatments. This will help clarify whether these species are unable to acclimate to external water conditions or if their capacity to do so is limited to a specific range.

### Asexual reproduction and invasiveness in urban area

Along with population growth, rapid urbanization has become a major feature of global change in recent decades, and urban areas are expected to continue expanding in the future (Li et al. 2021). Urban areas have unique climates due to the physical process of urbanization (Qian et al. 2022), such as generally drier atmosphere, light pollution from highly reflective buildings and artificial lighting at night, which are expected to influence plants in urban regions. As urbanization progresses, many native species struggling in adapting to urban ecosystem, while nonnative species with strong adaptive capacity are increasingly being introduced, with global ornamental horticulture being an important pathway (Dehnen-Schmutz et al. 2007, Hu et al. 2023).

In order to successfully invade and survive, plants must have specific combinations of traits that allow them to increase the population and expand the range rapidly, and to outcompete resident species in the nonnative ranges by gaining local dominance (Gioria et al. 2023). Regarding intrinsic factors, the asexual reproductive strategy of pseudoviviparous *Kalanchoe* species gave them the ability to reproduce throughout the year without limited by pollinators, which accelerated the process of recruitment (Piana et al. 2019). As for extrinsic factors, the climate and infertile land surface of urban ecosystems share similarities with the original habitat of these species.

Flexible adjustments in reproductive strategies to different environments was observed in this study, which may be strongly related to the invasive success of *Kalanchoe* pseudoviviparous plants in cities around the world (Fig. 4). *K. delagoensis* and *K*.*× houghtonii* produce significantly more plantlets in high light environment, while *K. daigremontiana* and *K. laetivirens* develop plantlets with significantly higher fresh weight. Both strategies are highly beneficial for plant recruitment. The species with smaller leaves seemed to perform better in the high light condition and as succulents they could outperform most plant species in the hot and dry area. With strong competitive advantages, increasing the plantlet number could contribute to faster colonization and population expansion. Species with larger leaves may suffer from the exposure to high radiation, but meanwhile can store more water in the leaf mesophyll. By putting more nutrition in each plantlet, the livability of offsprings is assured, as they will have carried abundant resources for independent growth and prosperity. In addition, the morphology of the plantlets matched that of the parent, helping them to thrive in the same environment, as dispersal distances are limited for such reproductive strategy.

The introduction of nonnative plants has both advantages and disadvantages. On one hand, there is the risk of plant invasion and its consequences, such as loss of local biodiversity; on the other hand, nonnative plants can provide additional ecosystem services. Many succulent species are used for green roofs, due to their ability to survive long periods of time between precipitation or irrigation events and have higher survival rate under extreme climatic conditions. These plants can also enhance the thermal insulation of buildings and alleviate the urban heat island effect (Vijayaraghavan 2016, Di Miceli et al. 2022). A sophisticated management and monitoring system for urban nonnative plants is currently needed, with particular attention to species with high invasive potential.

## Statements and declarations

## Author contributions

Yanhong Tang and Zhe Zhang conceived and designed the experiments. Material preparation and data collection were performed by Zhe Zhang. Data analysis and interpretation were performed by Zhe Zhang, Daisuke Sugiura and Wataru Yamori. The first draft of the manuscript was written by Zhe Zhang and all authors commented on previous versions of the manuscript. All authors read and approved the final manuscript.

## References

Alpert P, Simms EL (2002) The relative advantages of plasticity and fixity in different environments: when is it good for a plant to adjust? Evol Ecol 16:285–297

Bloom AJ, Chapin FS, Mooney HA (1985) Resource Limitation in Plants-An Economic Analogy. Annu Rev Ecol Syst 16:363–392

Borland AM, Barrera Zambrano VA, Ceusters J, Shorrock K (2011) The photosynthetic plasticity of crassulacean acid metabolism: an evolutionary innovation for sustainable productivity in a changing world. New Phytol 191:619–633

Dehnen-Schmutz K, Julia T, Perrings C, Williamson M (2007) The Horticultural Trade and Ornamental Plant Invasions in Britain. Conserv Biol 21:224–231

Di Miceli G, Iacuzzi N, Licata M, La Bella S, Tuttolomondo T, Aprile S (2022) Growth and development of succulent mixtures for extensive green roofs in a Mediterranean climate. PLoS One 17:e0269446

Dodd AN, Borland AM, Haslam RP, Griffiths H, Maxwell K (2002) Crassulacean acid metabolism: plastic, fantastic. J Exp Bot 53:569–580.

Eggli U (2003) Crassulaceae. In: Eggli U (ed) Illustrated Handbook of Succulent Plants: Crassulaceae. Springer Berlin Heidelberg, Berlin, Heidelberg. pp 5–374

Eggli U, Nyffeler R (2009) Living under temporarily arid conditions - succulence as an adaptive strategy. Bradleya 27:13–36

Elmqvist T, Cox PA (1996) The Evolution of Vivipary in Flowering Plants. Oikos 77:3–9

Franklin KA (2008) Shade avoidance. New Phytol 179:930–944

Garcês HMP, Champagne CEM, Townsley BT, Park S, Malhó R, Pedroso MC, Harada JJ, Sinha NR (2007) Evolution of asexual reproduction in leaves of the genus Kalanchoë. Proc Natl Acad Sci USA 104:15578–15583

Gideon FS (2020) Notes on Kalanchoe× Houghtonii (Crassulaceae Subfam. Kalanchooideae), an Early Hybrid between Two Species of K. Subg. Bryophyllum. Haseltonia 2019:78–85

Gioria M, Hulme PE, Richardson DM, Pyšek P (2023) Why Are Invasive Plants Successful? Annu Rev Plant Biol 74:635–670

Groner MG (1975) Allelopathic Influence of Kalanchoe daigremontiana on Other Species of Plants. Bot Gaz 136:207–211

Guerra-García A, Golubov J, Mandujano MC (2015) Invasion of Kalanchoe by clonal spread. Biol Invasions 17:1615–1622

Herrera I, Ferrer-Paris JR, Hernandez-Rosas JI, Nassar JM (2016) Impact of two invasive succulents on native-seedling recruitment in Neotropical arid environments. J Arid Environ 132:15–25

Herrera I, Hernandez M-J, Lampo M, Nassar JM (2012) Plantlet recruitment is the key demographic transition in invasion by Kalanchoe daigremontiana. Popul Ecol 54:225–237

Herrera I, Nassar JM (2009) Reproductive and recruitment traits as indicators of the invasive potential of Kalanchoe daigremontiana (Crassulaceae) and Stapelia gigantea (Apocynaceae) in a Neotropical arid zone. J Arid Environ 73:978–986

Howe MD (1931) A Morphological Study of the Leaf Notches of Bryophyllum Calycinum. Am J Bot 18:387–390

Hu S, Jin C, Liao R, Huang L, Zhou L, Long Y, Luo M, Jim CY, Hu W, Lin D, Chen S, Liu C, Jiang Y, Yang Y (2023) Herbaceous ornamental plants with conspicuous aesthetic traits contribute to plant invasion risk in subtropical urban parks. J Environ Manage 347:119059

Jacome-Blasquez F, Kim M (2023) Meristem genes are essential for the vegetative reproduction of Kalanchoe pinnata. Front Plant Sci 14:1157619

Li X, Zhou Y, Hejazi M, Wise M, Vernon C, Iyer G, Chen W (2021) Global urban growth between 1870 and 2100 from integrated high resolution mapped data and urban dynamic modeling. Commun Earth Environ 2:201

Li YC, Lin TC, Martin CE (2015) Leaf anthocyanin, photosynthetic light-use efficiency, and ecophysiology of the South African succulent Anacampseros rufescens (Anacampserotaceae). S Afr J Bot 99:122–128

Li Z, Ahammed GJ (2023) Plant stress response and adaptation via anthocyanins: A review. Plant Stress 10: 100230.

Lichtenthaler HK (1987) Chlorophylls and carotenoids: Pigments of photosynthetic biomembranes. Methods Enzymol 148: 350–382

Liscum E, Askinosie SK, Leuchtman DL, Morrow J, Willenburg KT, Coats DR (2014) Phototropism: Growing towards an Understanding of Plant Movement. The Plant Cell 26:38–55

Luttge U (2010) Ability of crassulacean acid metabolism plants to overcome interacting stresses in tropical environments. AoB Plants 2010:plq005

Matesanz S, Gianoli E, Valladares F (2010) Global change and the evolution of phenotypic plasticity in plants. Ann N Y Acad Sci 1206:35–55

Nicotra AB, Atkin OK, Bonser SP, Davidson AM, Finnegan EJ, Mathesius U, Poot P, Purugganan MD, Richards CL, Valladares F, van Kleunen M (2010) Plant phenotypic plasticity in a changing climate. Trends Plant Sci 15:684–692

Ogburn RM, Edwards EJ (2010) The Ecological Water-Use Strategies of Succulent Plants. In: Jean-Claude K, Michel D (ed) Advances in Botanical Research. Academic Press, pp 55:179–225

Ogburn RM, Edwards EJ (2012) Quantifying succulence: a rapid, physiologically meaningful metric of plant water storage. Plant Cell Environ 35:1533–1542

Perez-Lopez AV, Lim SD, Cushman JC (2023) Humboldt Review: Tissue succulence in plants: Carrying water for climate change. J Plant Physiol 289:154081

Piana MR, Aronson MF, Pickett ST, Handel SN (2019) Plants in the city: understanding recruitment dynamics in urban landscapes. Front Ecol Environ 17:455–463

Qian Y, Chakraborty TC, Li J, Li D, He C, Sarangi C, Chen F, Yang X, Leung LR (2022) Urbanization Impact on Regional Climate and Extreme Weather: Current Understanding, Uncertainties, and Future Research Directions. Adv Atmos Sci 39:819–860

Rojas-Sandoval J, Acevedo-Rodríguez P (2022a) Kalanchoe daigremontiana (devil’s backbone). CABI Compendium.

Rojas-Sandoval J, and Acevedo-Rodríguez P (2022b) Kalanchoe delagoensis (chandelier plant). CABI Compendium.

Smith GF, Figueiredo E, van Wyk AE (2019) Chapter 6 - Geographical Distribution and Ecology. In: Smith GF, Figueiredo E, van Wyk AE (ed) Kalanchoe (Crassulaceae) in Southern Africa. Academic Press, pp 45–54

Steyn WJ, Wand SJE, Holcroft DM, Jacobs G (2002) Anthocyanins in vegetative tissues: a proposed unified function in photoprotection. New Phytol 155:349–361

Stotz GC, Salgado-Luarte C, Escobedo VM, Valladares F, Gianoli E (2021) Global trends in phenotypic plasticity of plants. Ecol Lett 24:2267–2281

Valladares F, Martinez-Ferri E, Balaguer L, Perez-Corona E, Manrique E (2000) Low leaf-level response to light and nutrients in Mediterranean evergreen oaks: a conservative resource-use strategy? New Phytol 148:79–91

Vijayaraghavan K (2016) Green roofs: A critical review on the role of components, benefits, limitations and trends. Renewable Sustainable Energy Rev 57:740–752

Wang ZQ, Guillot D, Ren MX, López-Pujol J (2016) Kalanchoe (Crassulaceae) as invasive aliens in China – new records, and actual and potential distribution. Nord J Bot 34:349–354

Weiner J (2004) Allocation, plasticity and allometry in plants. Perspect Plant Ecol Evol Syst 6:207–215

Xu H, Qiang S, Genovesi P, Ding H, Wu J, Meng L, Han Z, Miao J, Hu B, Guo J, Sun H, Huang C, Lei J, Le Z, Zhang X, He S, Wu Y, Zheng Z, Chen L, Jarošik V, Pysek P (2012) An inventory of invasive alien species in China. NeoBiota 15:1–26

