## Supplemental Figures and Tables for "Morphological Plasticity and Reproductive Strategies of *Kalanchoe* Species in Invasive Spread"

Supplemental Fig. 1

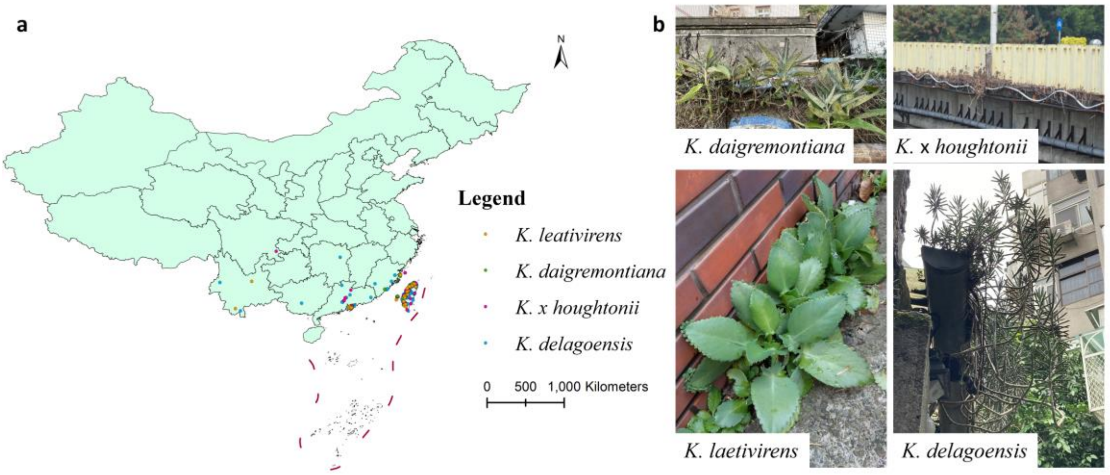


**Fig. S1** Distribution of four *Kalanchoe* pseudoviviparous species in China. (a) Distribution map of four species in China, mainly on the north of China. Occurrence data gathered from iNaturalist, the Global Biodiversity Information Facility (GBIF), the Plant Photo Bank of China (PPBC), the Chinese Field Herbarium (CFH) and the Chinese Virtual Herbarium (CVH). (b) Pictures of four species in the urban areas of China, acquired from iNaturalist

Supplemental Fig. 2

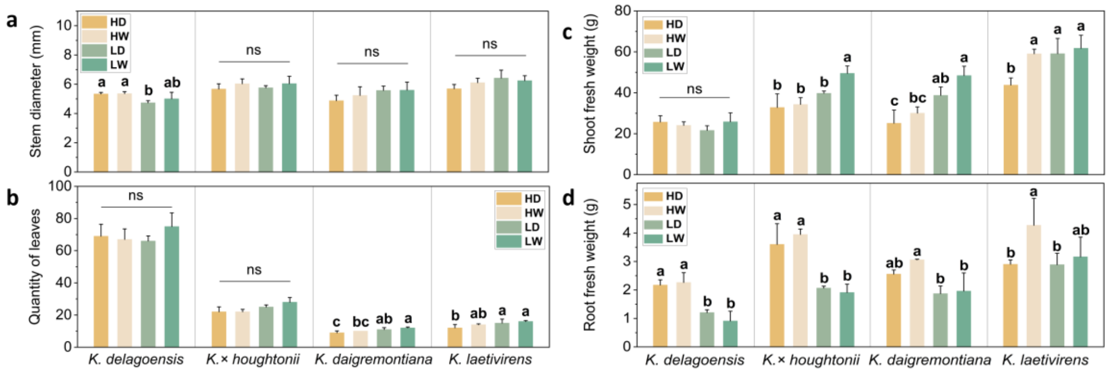


**Fig. S2** Individual morphology and biomass allocation of four *Kalanchoe* species under various environmental conditions. Showing (a) stem diameter, (b) quantity of leaves, (c) shoot fresh weight and (d) root fresh weight. Difference of alphabet above the error bar shows significance, while *ns* means no significant difference between the treatments

Supplemental Fig. 3

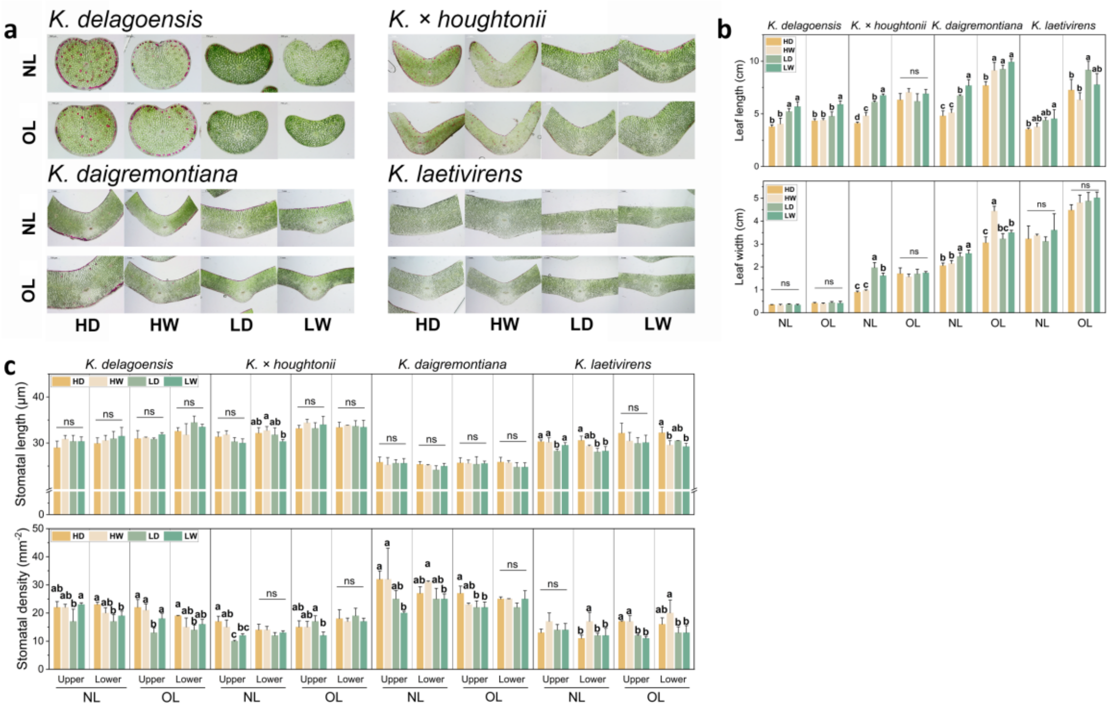


**Fig. S3** Leaf morphological traits of four *Kalanchoe* species under various environmental conditions. NL represents new leaves and OL represents old leaves. (a) Cross section of leaves middle from plants grown in various environmental conditions observed under microscope. (b) Bar graphs of leaf morphological traits. (c) Bar graphs of leaf stomatal length and density. Difference of alphabet above the error bar shows significance, while *ns* means no significant difference between the treatments

Supplemental Table 1

**Table S1** Asexual reproduction traits of each species under various light and water environments

| Species | Treatment | Fresh weight of single plantlet | Quantity of plantlets | Plantlet total mass | Reproduction ratio |
|---|---|---|---|---|---|
| *K. delagoensis*is | HD | 0.015 ± 0.002 c | 204 ± 47 a | 3.115 ± 0.954 a | 0.111 ± 0.024 b |
| | HW | 0.014 ± 0.001 c | 253 ± 18 a | 3.465 ± 0.140 a | 0.132 ± 0.011 b |
| | LD | 0.043 ± 0.002 a | 118 ± 54 b | 4.988 ± 2.196 a | 0.214 ± 0.078 a |
| | LW | 0.030 ± 0.004 b | 114 ± 25 b | 3.455 ± 1.221 a | 0.128 ± 0.027 b |
| *K.* x *houghtonii* | HD | 0.025 ± 0.016 a | 183 ± 6 a | 4.649 ± 0.821 a | 0.130 ± 0.023 ab |
| | HW | 0.029 ± 0.005 a | 187 ± 12 a | 5.385 ± 0.948 a | 0.143 ± 0.035 a |
| | LD | 0.036 ± 0.010 a | 142 ± 16 b | 5.004 ± 1.107 a | 0.120 ± 0.028 ab |
| | LW | 0.034 ± 0.005 a | 127 ± 10 b | 4.327 ± 0.841 a | 0.084 ± 0.010 b |
| *K. daigremontiana* | HD | 0.037 ± 0.005 a | 123 ± 20 a | 4.562 ± 1.006 a | 0.170 ± 0.048 a |
| | HW | 0.031 ± 0.005 a | 167 ± 10 a | 5.127 ± 0.477 a | 0.156 ± 0.029 a |
| | LD | 0.021 ± 0.004 b | 117 ± 19 a | 2.534 ± 0.806 b | 0.063 ± 0.022 b |
| | LW | 0.022 ± 0.001 b | 122 ± 42 a | 2.705 ± 0.810 b | 0.055 ± 0.019 b |
| *K. laetivirens* | HD | 0.134 ± 0.022 a | 83 ± 36 a | 10.621 ± 2.661 a | 0.226 ± 0.044 a |
| | HW | 0.104 ± 0.013 a | 130 ± 8 a | 13.575 ± 2.480 a | 0.214 ± 0.033 a |
| | LD | 0.061 ± 0.011 b | 81±34 a | 4.915 ± 2.167 b | 0.079 ± 0.030 b |
| | LW | 0.046 ± 0.018 b | 92±37 a | 4.339 ± 2.969 b | 0.070 ± 0.054 b |

Supplemental Fig. 4

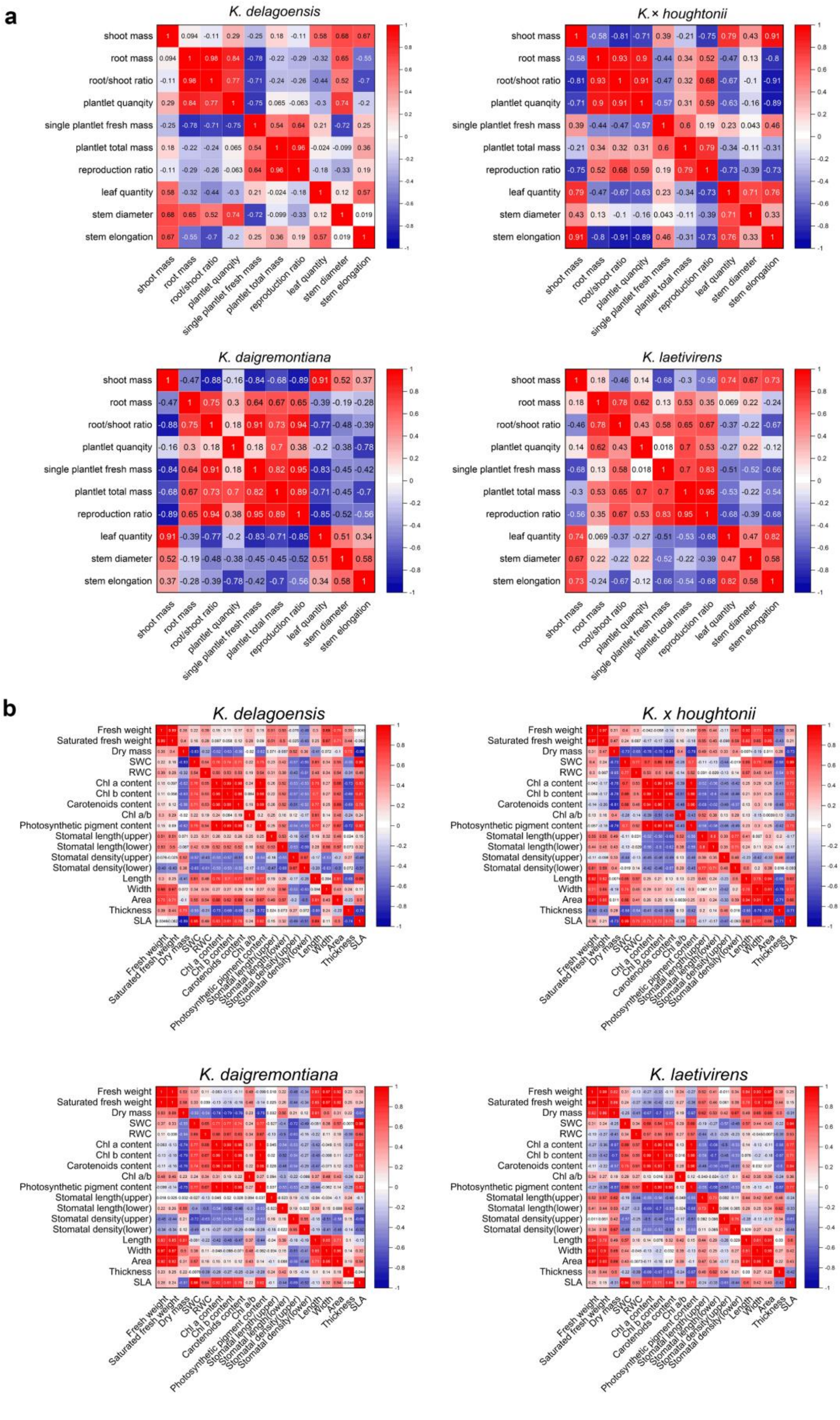


**Fig. S4** Correlation heatmap of (a) individual traits and (b) leaf traits of each species

Supplemental Table 2

**Table S2** Result of two-way ANOVA about the effects of light, water and their interactions on plant leaf traits

| Traits | Species | Leaf | F-value and significance | | |
|---|---|---|---|---|---|
| | | Type | Light | Water | Light*Water |
| Leaf length | *K. delagoensis* | NL | 46.626*** | 2.688 | 0.215 |
| | | OL | 33.805*** | 12.128** | 9.872* |
| | *K.* x *houghtonii* | NL | 344.055*** | 38.668*** | 0.079 |
| | | OL | 0.183 | 5.227 | 0.000 |
| | *K. daigremontiana* | NL | 81.765*** | 6.913* | 1.68 |
| | | OL | 25.499*** | 14.893** | 1.975 |
| | *K. laetivirens* | NL | 8.755* | 0.433 | 0.031 |
| | | OL | 10.311* | 4.948 | 0.207 |
| Leaf width | *K. delagoensis* | NL | 0.965 | 2.874 | 0.493 |
| | | OL | 0.837 | 0.644 | 0.023 |
| | *K.* x *houghtonii* | NL | 133.048*** | 3.361 | 8.171* |
| | | OL | 0.702 | 0.226 | 0.747 |
| | *K. daigremontiana* | NL | 25.212** | 1.665 | 0.084 |
| | | OL | 3.84 | 36.84*** | 19.25** |
| | *K. laetivirens* | NL | 0.068 | 1.378 | 0.43 |
| | | OL | 2.998 | 1.826 | 0.295 |
| Leaf thickness | *K. delagoensis* | NL | 102.771*** | 2.199 | 3.504 |
| | | OL | 50.268*** | 6.011* | 0.191 |
| | *K.* x *houghtonii* | NL | 84.078*** | 1.218 | 9.923* |
| | | OL | 1.098 | 1.659 | 2.776 |
| | *K. daigremontiana* | NL | 9.974* | 2.181 | 0 |
| | | OL | 0.162 | 1.399 | 0.256 |
| | *K. laetivirens* | NL | 10.95* | 0.000 | 5.31 |
| | | OL | 5.532* | 2.921 | 1.332 |
| Leaf area | *K. delagoensis* | NL | 14.490** | 0.766 | 0.001 |
| | | OL | 7.836* | 4.357 | 4.977 |
| | *K.* x *houghtonii* | NL | 439.602*** | 2.353 | 1.995 |
| | | OL | 0.06 | 0.999 | 0.184 |
| | *K. daigremontiana* | NL | 41.634*** | 10.847* | 0.743 |
| | | OL | 3.773 | 28.015** | 8.078* |
| | *K. laetivirens* | NL | 1.241 | 0.628 | 0.386 |
| | | OL | 4.094 | 0.186 | 0.134 |
| SLA | *K. delagoensis* | NL | 224.44*** | 4.000 | 1.468 |
| | | OL | 11.858** | 0.027 | 1.474 |
| | *K.* x *houghtonii* | NL | 830.588*** | 1.206 | 4.901 |
| | | OL | 65.089*** | 0.09 | 0.054 |
| | *K. daigremontiana* | NL | 42.254*** | 4.415 | 0.410 |
| | | OL | 111.265*** | 3.183 | 0.000 |
| | *K. laetivirens* | NL | 86.517*** | 0.099 | 0.575 |
| | | OL | 27.004*** | 0.283 | 0.415 |

| | | | | | |
|---|---|---|---|---|---|
| Stomata length (upper) | *K. delagoensis* | NL | 0.43 | 1.889 | 1.913 |
| | | OL | 0.272 | 1.327 | 0.559 |
| | *K.* x *houghtonii* | NL | 7.242* | 0.02 | 0.485 |
| | | OL | 0.071 | 2.02 | 0.096 |
| | *K. daigremontiana* | NL | 0.001 | 0.141 | 0.163 |
| | | OL | 0.069 | 0.005 | 0.031 |
| | *K. laetivirens* | NL | 13.605** | 2.414 | 3.479 |
| | | OL | 1.629 | 0.574 | 0.845 |
| Stomata length (lower) | *K. delagoensis* | NL | 1.316 | 0.471 | 0.003 |
| | | OL | 4.41 | 1.017 | 0.013 |
| | *K.* x *houghtonii* | NL | 4.644 | 0.507 | 2.515 |
| | | OL | 0.011 | 0.037 | 0.251 |
| | *K. daigremontiana* | NL | 3.077 | 0.797 | 1.522 |
| | | OL | 3.42 | 0.015 | 0.018 |
| | *K. laetivirens* | NL | 15.586** | 1.304 | 2.524 |
| | | OL | 4.214 | 12.963** | 1.490 |
| Stomatal density (upper) | *K. delagoensis* | NL | 1.44 | 3.44 | 2.679 |
| | | OL | 20.417** | 2.817 | 4.817 |
| | *K.* x *houghtonii* | NL | 30.000*** | 0.133 | 6.533* |
| | | OL | 0.022 | 3.756 | 5 |
| | *K. daigremontiana* | NL | 10.351* | 0.791 | 0.633 |
| | | OL | 9.436* | 2.118 | 1.796 |
| | *K. laetivirens* | NL | 0.417 | 2.817 | 0.817 |
| | | OL | 53.895*** | 0.842 | 0.211 |
| Stomatal density (lower) | *K. delagoensis* | NL | 11.111* | 0.111 | 7.111* |
| | | OL | 3.25 | 0.481 | 5.558* |
| | *K.* x *houghtonii* | NL | 4.05 | 0.45 | 0.05 |
| | | OL | 0.138 | 0.754 | 0.015 |
| | *K. daigremontiana* | NL | 6.280* | 0.824 | 0.61 |
| | | OL | 1.696 | 1.254 | 1.003 |
| | *K. laetivirens* | NL | 2.571 | 4.571 | 5.786* |
| | | OL | 7.577* | 1.523 | 1.09 |
| Leaf fresh weight | *K. delagoensis* | NL | 1.547 | 0.193 | 0.46 |
| | | OL | 1.657 | 1.949 | 2.036 |
| | *K.* x *houghtonii* | NL | 152.427*** | 11.664** | 2.289 |
| | | OL | 1.396 | 3.011 | 0.177 |
| | *K. daigremontiana* | NL | 12.093* | 3.951 | 3.186 |
| | | OL | 3.283 | 40.115*** | 11.073* |
| | *K. laetivirens* | NL | 0.107 | 0.583 | 1.243 |
| | | OL | 0.12 | 0.216 | 0.066 |
| Leaf saturated Fresh weight | *K. delagoensis* | NL | 0.841 | 0.14 | 0.295 |
| | | OL | 3.061 | 1.243 | 3.061 |
| | *K.* x *houghtonii* | NL | 79.452*** | 6.848* | 0.890 |
| | | OL | 8.829* | 2.495 | 0.116 |

| | | | | | |
|---|---|---|---|---|---|
| | *K. daigremontiana* | NL | 10.712* | 3.788 | 3.495 |
| | | OL | 5.201 | 31.523*** | 10.826* |
| | *K. laetivirens* | NL | 0.86 | 0.336 | 1.103 |
| | | OL | 0.192 | 0.235 | 0.094 |
| Leaf dry weight | *K. delagoensis* | NL | 38.009*** | 0.147 | 0.026 |
| | | OL | 8.327* | 0.001 | 3.601 |
| | *K.* x *houghtonii* | NL | 149.670*** | 5.466* | 0.275 |
| | | OL | 145.670*** | 0.882 | 0.027 |
| | *K. daigremontiana* | NL | 27.792** | 1.272 | 1.166 |
| | | OL | 88.248*** | 3.281 | 2.465 |
| | *K. laetivirens* | NL | 23.106** | 2.028 | 0.415 |
| | | OL | 3.949 | 0.252 | 0.001 |
| SWC | *K. delagoensis* | NL | 254.293*** | 2.515 | 0.031 |
| | | OL | 4.458 | 0.061 | 1.368 |
| | *K.* x *houghtonii* | NL | 335.500*** | 0.027 | 1.346 |
| | | OL | 97.592*** | 0.531 | 0.076 |
| | *K. daigremontiana* | NL | 31.551*** | 3.343 | 0.849 |
| | | OL | 78.458*** | 7.130* | 0.129 |
| | *K. laetivirens* | NL | 32.489*** | 0.306 | 1.79 |
| | | OL | 10.802* | 0.038 | 0.246 |
| RWC | *K. delagoensis* | NL | 6.267* | 0.67 | 1.644 |
| | | OL | 3.61 | 2.44 | 1.291 |
| | *K.* x *houghtonii* | NL | 63.017*** | 3.527 | 5.866* |
| | | OL | 38.420*** | 0.764 | 0.38 |
| | *K. daigremontiana* | NL | 10.683* | 1.081 | 0.009 |
| | | OL | 8.425* | 9.275* | 0.025 |
| | *K. laetivirens* | NL | 2.538 | 0.159 | 0.162 |
| | | OL | 6.408* | 0.466 | 0.003 |
| Chl a content | *K. delagoensis* | NL | 207.977*** | 9.518* | 1.522 |
| | | OL | 622.333*** | 0.006 | 2.849 |
| | *K.* x *houghtonii* | NL | 41.472*** | 1.672 | 9.267* |
| | | OL | 40.065*** | 1.665 | 0.331 |
| | *K. daigremontiana* | NL | 237.80*** | 41.25*** | 0.01 |
| | | OL | 21.053** | 1.164 | 0.317 |
| | *K. laetivirens* | NL | 65.922*** | 3.158 | 0.392 |
| | | OL | 61.710*** | 0.418 | 0.29 |
| Chl b content | *K. delagoensis* | NL | 94.72*** | 3.6 | 0.05 |
| | | OL | 279.146*** | 3.4 | 8.854* |
| | *K.* x *houghtonii* | NL | 37.751*** | 0.67 | 4.799 |
| | | OL | 24.018** | 0.688 | 0.127 |
| | *K. daigremontiana* | NL | 278.003*** | 39.959*** | 1.032 |
| | | OL | 38.747*** | 0.763 | 0.955 |
| | *K. laetivirens* | NL | 72.305*** | 5.684* | 1.324 |
| | | OL | 70.420*** | 0.202 | 0.249 |

| | | | | | |
|---|---|---|---|---|---|
| Carotenoids content | *K. delagoensis* | NL | 61.573*** | 1.645 | 0.07 |
| | | OL | 122.185*** | 0.005 | 0.865 |
| | *K.* x *houghtonii* | NL | 59.358*** | 0.091 | 4.252 |
| | | OL | 22.983** | 0.336 | 0.35 |
| | *K. daigremontiana* | NL | 83.298*** | 10.822* | 2.973 |
| | | OL | 45.654*** | 0.748 | 0 |
| | *K. laetivirens* | NL | 39.080*** | 1.738 | 0.038 |
| | | OL | 81.411*** | 0.047 | 2.889 |
| Chl a/b | *K. delagoensis* | NL | 2.352 | 0.001 | 1.256 |
| | | OL | 0.082 | 6.628* | 5.705* |
| | *K.* x *houghtonii* | NL | 3.152 | 0.412 | 1.922 |
| | | OL | 0.279 | 0.441 | 0.112 |
| | *K. daigremontiana* | NL | 2.238 | 0.929 | 0.001 |
| | | OL | 0.297 | 5.062 | 3.915 |
| | *K. laetivirens* | NL | 2.242 | 2.633 | 3.252 |
| | | OL | 0.049 | 0.624 | 0.154 |
| Total photosynthetic pigment contents | *K. delagoensis* | NL | 160.149*** | 6.747* | 0.598 |
| | | OL | 432.574*** | 0.262 | 3.928 |
| | *K.* x *houghtonii* | NL | 43.039*** | 1.21 | 7.707* |
| | | OL | 36.680*** | 1.298 | 0.163 |
| | *K. daigremontiana* | NL | 259.150*** | 41.781*** | 0.248 |
| | | OL | 26.543** | 1.06 | 0.363 |
| | *K. laetivirens* | NL | 66.842*** | 3.632 | 0.506 |
| | | OL | 69.021*** | 0.322 | 0.456 |

***: $p<0.01$, **: $p<0.05$, *: $p<0.1$

Supplemental Table 3

**Table S3** Result of two-way ANOVA about the effects of light, water and their interactions on plant individual traits

| Traits | Species | F-value and significance | | |
|---|---|---|---|---|
| | | Light | Water | Light*Water |
| Shoot fresh weight | *K. delagoensis* | 0.451 | 0.56 | 2.85 |
| | *K.* x *houghtonii* | 21.136** | 5.437* | 2.953 |
| | *K. daigremontiana* | 31.163*** | 6.499* | 0.635 |
| | *K. laetivirens* | 8.751* | 8.583* | 4.251 |
| Root fresh weight | *K. delagoensis* | 57.648*** | 0.45 | 1.588 |
| | *K.* x *houghtonii* | 58.627*** | 0.167 | 1.158 |
| | *K. daigremontiana* | 13.561** | 1.403 | 0.797 |
| | *K. laetivirens* | 2.417 | 5.222 | 2.309 |
| Root/shoot ratio | *K. delagoensis* | 55.509*** | 0.952 | 6.536* |
| | *K.* x *houghtonii* | 304.438*** | 0.978 | 6.305* |
| | *K. daigremontiana* | 27.638** | 0.348 | 0.023 |
| | *K. laetivirens* | 7.540* | 0.402 | 0.038 |

| | | | | |
|---|---|---|---|---|
| Quantity of leaves | *K. delagoensis* | 0.555 | 0.938 | 2.936 |
| | *K.* x *houghtonii* | 10.361** | 2.288 | 0.935 |
| | *K. daigremontiana* | 19.861** | 7.013* | 0.000 |
| | *K. laetivirens* | 6.342* | 2.675 | 0.405 |
| Stem diameter | *K. delagoensis* | 9.011* | 0.997 | 0.604 |
| | *K.* x *houghtonii* | 0.051 | 2.453 | 0.033 |
| | *K. daigremontiana* | 3.832 | 0.493 | 0.385 |
| | *K. laetivirens* | 3.879 | 0.282 | 1.784 |
| Stem elongation | *K. delagoensis* | 10.667* | 10.293* | 1.535 |
| | *K.* x *houghtonii* | 264.86*** | 56.68*** | 31.40*** |
| | *K. daigremontiana* | 1.58 | 0.056 | 0.12 |
| | *K. laetivirens* | 14.868** | 1.011 | 0.257 |
| Fresh weight of single plantlet | *K. delagoensis* | 268.14*** | 26.27*** | 19.14** |
| | *K.* x *houghtonii* | 4.109 | 0.07 | 0.497 |
| | *K. daigremontiana* | 27.948** | 0.757 | 2.054 |
| | *K. laetivirens* | 46.906*** | 5.646 | 0.703 |
| Quantity of plantlets | *K. delagoensis* | 25.092** | 1.028 | 1.384 |
| | *K.* x *houghtonii* | 57.452*** | 0.601 | 2.178 |
| | *K. daigremontiana* | 1.712 | 1.9 | 1.383 |
| | *K. laetivirens* | 1.224 | 2.573 | 1.028 |
| Total fresh weight of plantlets | *K. delagoensis* | 1.436 | 0.58 | 1.468 |
| | *K.* x *houghtonii* | 0.423 | 0.003 | 1.711 |
| | *K. daigremontiana* | 18.473** | 0.467 | 0.150 |
| | *K. laetivirens* | 25.047** | 0.635 | 1.399 |
| Reproduction ratio | *K. delagoensis* | 3.935 | 1.731 | 4.654 |
| | *K.* x *houghtonii* | 5.304 | 0.556 | 2.748 |
| | *K. daigremontiana* | 29.916*** | 0.306 | 0.020 |
| | *K. laetivirens* | 37.537*** | 0.194 | 0.006 |

***: $p$<0.01, **: $p$<0.05, *: $p$<0.1
